# Run-length compressed metagenomic read classification with SMEM-finding and tagging

**DOI:** 10.1101/2025.02.25.640119

**Authors:** Lore Depuydt, Omar Y. Ahmed, Jan Fostier, Ben Langmead, Travis Gagie

## Abstract

Metagenomic read classification is a fundamental task in computational biology, yet it remains challenging due to the scale, diversity, and complexity of sequencing datasets. We propose a novel, run-length compressed index based on the move structure that enables efficient multi-class metagenomic classification in *O*(*r*) space, where *r* is the number of character runs in the BWT of the reference text. Our method identifies all super-maximal exact matches (SMEMs) of length at least *L* between a read and the reference dataset and associates each SMEM with one class identifier using a sampled tag array. A consensus algorithm then compacts these SMEMs with their class identifier into a single classification per read. We are the first to perform run-length compressed read classification based on full SMEMs instead of semi-SMEMs. We evaluate our approach on both long and short reads in two conceptually distinct datasets: a large bacterial pan-genome with few metagenomic classes and a smaller 16S rRNA gene database spanning thousands of genera or classes. Our method consistently outperforms SPUMONI 2 in accuracy and runtime while maintaining the same asymptotic memory complexity of *O*(*r*). Compared to Cliffy, we demonstrate better memory efficiency while achieving superior accuracy on the simpler dataset and comparable performance on the more complex one. Overall, our implementation carefully balances accuracy, runtime, and memory usage, offering a versatile solution for metagenomic classification across diverse datasets. The open-source C++11 implementation is available at https://github.com/biointec/tagger under the AGPL-3.0 license.

## 1 Introduction

Metagenomic read classification, where reads of unknown origin are matched against a reference set of candidate genomes or genes, has been a long-standing challenge in bioinformatics due to computational complexity, large and incomplete reference datasets, and the inherent noisiness of sequencing reads. Reads can be classified at different taxonomic levels—such as strain, species, or genus—depending on the question at hand. Metagenomic analyses are revolutionizing diverse fields, including infectious disease diagnostics [1], cancer prediction [2], antibiotic resistance surveillance [3], ecology research [4], and sustainable agriculture [5]. Many established metagenomic classification tools exist, such as Kraken 2 [6], Centrifuge [7], and MetaPhlAn [8, 9], each employing distinct approaches to classification. Kraken 2 uses a probabilistic, compact hash table that maps minimizers onto the lowest common ancestor (LCA) taxa [6]. Centrifuge employs an FM-index [10], based on the Burrows-Wheeler transform (BWT) [11], to find exact matches between reads and candidate references [7]. MetaPhlAn(4) relies on a preprocessed subset of clade-specific marker genes for classification [8, 9]. However, as reference databases continue to grow, these tools face scalability challenges. Lossless tools such as Centrifuge experience index growth proportional to the sequence content in the reference, which becomes impractical even for high-RAM workstations. Lossy tools such as Kraken 2 and MetaPhlAn lose specificity as database size increases [12]. More specifically, for *k*-mer-based tools like Kraken 2, there is no single value of *k* that consistently yields optimal results across all scenarios [12, 13].

Orthogonally, recent advancements have introduced a new wave of bioinformatics tools leveraging the compressibility of the BWT to create compact and efficient indexes, particularly for large, repetitive datasets. This approach was first applied in the run-length FM-index (RLFM-index) [14] and the r-index [15, 16], both achieving *O*(*r*) space complexity, where *r* is the number of character runs in the BWT. The introduction of the move structure [17] further accelerated this functionality by supporting *O*(1)-time core LF-operations while maintaining *O*(*r*) memory complexity. Building on these foundations, alignment tools such as MONI [18], PHONI [19], SPUMONI [20], Movi [21], and b-move [22, 23] have demonstrated the practical utility of run-length compressed indexes. These tools focus on tasks such as computing matching statistics or pseudo-matching lengths, finding and extending maximal exact matches, or lossless approximate pattern matching. In metagenomic classification, tools like SPUMONI 2 [24], Centrifuger [25], and Cliffy [26] have also applied run-length compression to create memory-efficient solutions. SPUMONI 2 performs multi-class metagenomic classification in *O*(*r*) space using a sampled document array and pseudomatching lengths, based on MONI’s matching statistics [24, 18]. Optional lossy minimizer digestion can further accelerate and compress the process with minimal accuracy penalties. Centrifuger supports taxonomic classification, i.e., mapping reads to the LCA of their matches in the taxonomic tree. It uses a lossless run-block compressed BWT of the reference to find semi-maximal matches between reads and references [25], though its worst-case space complexity remains *O*(*n*). Cliffy, though mainly intended for taxonomic classification, also supports multi-class metagenomic classification (referred to as document listing) using a run-length compressed BWT with document listing profiles [27]. It achieves an *O*(*rd*) space complexity, where *d* is the number of classes. Optionally, Cliffy’s lossy cliff compression significantly reduces index size while still supporting *approximate* document listing and accurate taxonomic classification [26].

Though these tools demonstrate the potential of run-length compressed indexes in metagenomic classification, our experiments show that there is still a need for further improvements in balancing space efficiency, scalability, performance, and classification accuracy. One limitation is that these tools build upon the r-index’s pattern matching principles, which can suffer significant computational overhead due to the multitude of cache misses introduced by complex memory access patterns [22]. Moreover, they all perform backward search to find semi-maximal matches but lack support for *full* maximal exact match-based classification. By considering only semi-maximal matches, specificity is reduced, which is particularly critical for distinguishing closely related species or even strains lower in the phylogenetic tree. In contrast, full maximal exact matches ensure that all reported matches are truly non-extendable, reducing ambiguity in classification.

### Contributions

We propose a new, run-length compressed index that efficiently supports multi-class metagenomic classification in *O*(*r*) space. Classification begins by identifying all *full* super-maximal exact matches (SMEMs) of a read relative to the reference dataset, where an SMEM is defined as an exact match that cannot be extended in either direction of the read while still occurring in the reference dataset [28].

The ability to compute all SMEMs encompasses all other exact matching queries, including various forms of half-maximal match queries and *k*-mer searches, as these can all be derived from the full set of SMEMs. To improve specificity, only SMEMs of length at least *L* are considered, found efficiently using a method similar to that described by Gagie et al. [29] (see also [30]).

During SMEM identification, we use a sampled tag array of class identifiers [31] to track the identifier of *one* metagenomic class in which the SMEM occurs. To ensure *O*(*r*) space complexity, the tag array is sampled at positions marking the end of a BWT run. After identifying SMEMs and their associated class identifiers, a consensus algorithm aggregates the results into a single classification per read. In addition to these theoretical advancements in memory usage and accuracy, we leverage a move structure-based index for high performance, irrespective of theoretical time complexities.

We evaluate our approach in terms of accuracy, runtime, and memory usage by comparing it to SPUMONI 2 and Cliffy, and observe that our approach achieves the best balance across all metrics. Its versatility is demonstrated by classifying both long and short reads on two conceptually different datasets.

## 2 Methods

### 2.1 Preliminaries

In this paper, the search text *T*, with length *n* = |*T*|, represents the concatenation of all candidate sequences for classification. *T* is expected to be a string over the alphabet ∑ ={ A, C, G, T}. If a candidate sequence contains a non-ACGT character, it is replaced with the dummy character ‘X’, which is never matched to ensure that no SMEMs span such characters. Additionally, the same dummy character ‘X’ is placed between each candidate sequence to prevent SMEMs from spanning multiple sequences. The concatenated text *T* is always terminated with the sentinel character ‘$’, which is lexicographically smaller than any other character in the extended alphabet ∑^*′*^ = {$, A, C, G, T, X}

Furthermore, we use the following notation. Array and string indexing start at 0. A substring within a text *T* is defined by a half-open interval [*i, j*) over *T*, where 0≤ *i* ≤*j* ≤*n*. An empty interval is denoted by [*i, i*). Intervals over other data structures, like the BWT, will also be presented as half-open intervals.

#### 2.1.1 Backward and forward character matching with move tables

Finding *full* SMEMs of a read relative to the candidate dataset requires supporting backward and forward character extensions in the index. To achieve this, we leverage the cache-efficient principles of the move structure [17]. Our implementation follows the approach introduced for b-move. In this section, we briefly recapitulate the elements necessary for SMEM-finding in the context of this paper. For a more detailed overview, including pseudo-code, we refer the reader to the b-move papers [22, 23].

In the move structure, a move table facilitates fast LF-operations by mapping a character in the Last column of the lexicographically ordered rotations of *T* (i.e., the BWT) to its corresponding position in the First column. The LF move table *M* for a search text *T* consists of *r* rows, each representing a run in the BWT. Each row consists of four values: the run character *c*, the BWT start index *p* of the run, the LF mapping *π* = LF(*p*), and the run index *ξ* that contains BWT index *π*. The left panel of Table 1 shows the move table *M* for a small conceptual example consisting of two strains: “CTATGTC” and “ATATGTTGGTC”. For completeness, the corresponding FM-index is provided in Supplementary Table S1.

**Table 1:**
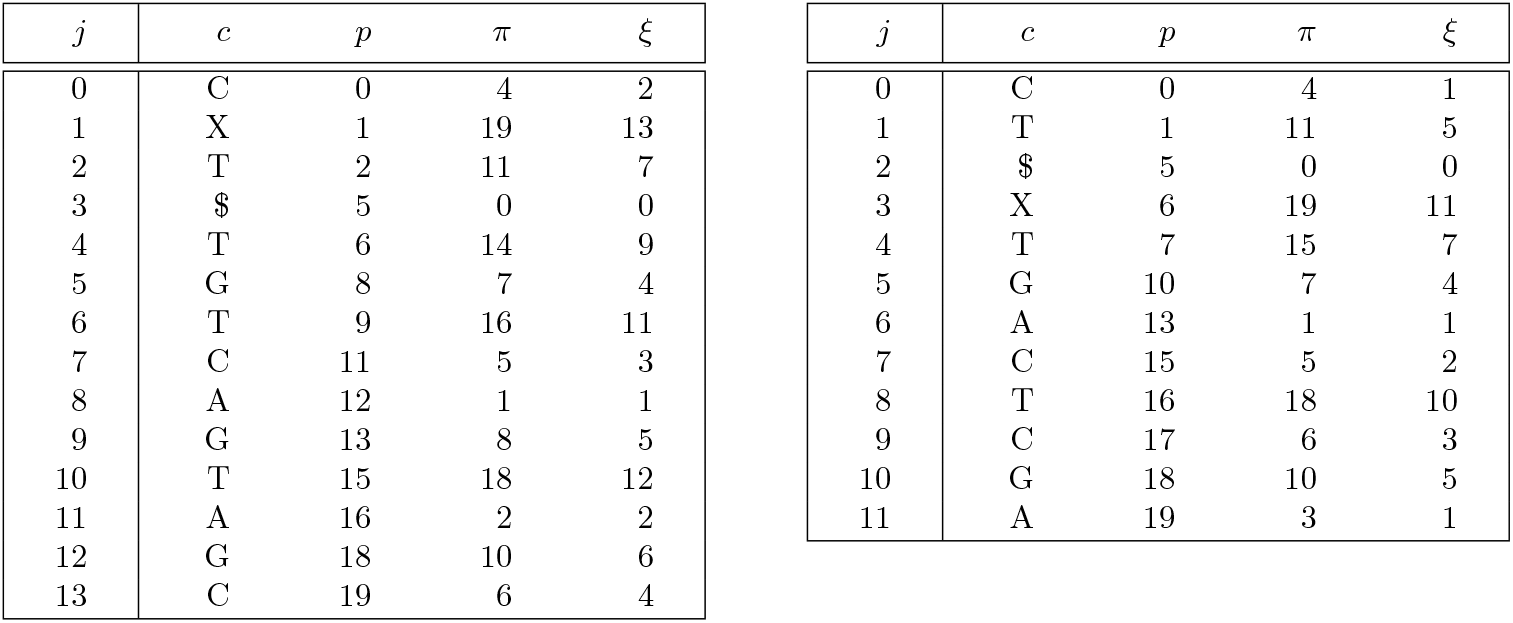
Left: Move table *M* for the search text *T* = “CTATGTCXATATGTTGGTC$”. Right: Move table *M* ^rev^ for the reverse search text *T* ^rev^ = “$CTGGTTGTATAXCTGTATC”. The corresponding FM-indexes are provided in Supplementary Tables S1 and S2.

Since all BWT positions within the same run map to consecutive positions in the First column, the move table *M* suffices to perform the LF operation by accessing only a single row. Specifically, the LF mapping of BWT index *i*, located in run *j*, is calculated as *i*^*′*^ = *M* [*j*].*π* + (*i* −*M* [*j*].*p*). To determine the run where *i*^*′*^ resides, we initially check whether run *M* [*j*].*ξ* contains index *i*^*′*^. If not, we fast forward through subsequent runs until the correct run *j*^*′*^ containing index *i*^*′*^ is located. This functionality is bundled in the function *M*.`LF`(*i, j*) = (*i*^*′*^, *j*^*′*^), which returns the LF mapping of a BWT index and its corresponding run.

Using this LF functionality, we can perform backward character extensions with the move table as follows. Let the interval [*s, e*) over the BWT represent a pattern *P*, meaning all lexicographically sorted rotations of *T* within this interval are prefixed by *P*. The indices *s* and *e* − 1 lie within runs *R*_*s*_ and *R*_*e*_, respectively. To extend *P* to *cP*, we must identify the smallest interval [*s*_*c*_, *e*_*c*_) ⊆ [*s, e*) that contains all occurrences of character *c* in the BWT within [*s, e*), along with their updated run indices *R*_*s,c*_ and *R*_*e,c*_. This can be achieved by traversing inward over the rows of the move table, starting from runs *R*_*s*_ and *R*_*e*_. This functionality is encapsulated in the functions *M*.`walkToNextRun`([*s, e*), *R*_*s*_, *R*_*e*_, *c*) (returning (*s*_*c*_, *R*_*s,c*_)) and *M*.`walkToPreviousRun`([*s, e*), *R*_*s*_, *R*_*e*_, *c*) (returning (*e*_*c*_, *R*_*e,c*_)). Once [*s*_*c*_, *e*_*c*_) and the corresponding run indices are found, we compute the updated BWT interval [*s*^*′*^, *e*^*′*^) for pattern *cP*, where *s*^*′*^ and *e*^*′*^ − 1 are located in runs *Rs′* and *Re′*, respectively. The output is computed as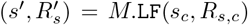and (*e*^*′*^ − 1, *Re′*) = *M*.LF(*e*_*c*_ − 1, *R*_*e,c*_). This entire process is encapsulated in the function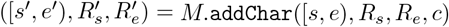.

#### Example

Assume we want to backward match the pattern “TGT” using the move table *M* on the left in Table 1. We initialize the empty move interval as ([0, 20), 0, 13), spanning all positions in the BWT and all rows in *M*. To match the last character “T”, we first call *M*.`walkToNextRun` and *M*.`walkToPreviousRun` to find the first and last occurrences of “T” in the BWT along with their corresponding runs. This results in pairs (2, 2) and (15, 10), respectively. Performing the LF mapping on these (position, run) pairs gives (11, 7) and (18, 12). Thus, the move interval for “T” is ([11, 19), 7, 12). Similarly, two subsequent calls to *M*.`addChar` for characters “G” and “T” yield move intervals ([8, 11), 5, 6) and ([16, 18), 11, 11), respectively.

The overview above describes backward character matching using move table *M*, enabling the extension of *P* to *cP*. However, our full SMEM-finding algorithm also requires support for forward matching, i.e., extending *P* to *Pc*. To facilitate this symmetrical functionality, we store a second move table, *M* ^rev^, which represents the reverse of the search text, *T* ^rev^, in *r*^rev^ rows. Using *M* ^rev^, pattern *P* can be extended to *Pc* by computing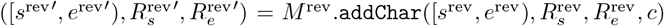. The right panel of Table 1 shows the move table *M* ^rev^, corresponding to our two example strains. For completeness, the FM-index for the reverse search text is provided in Supplementary Table S2.

Theoretically, function `addChar` has a worst-case time complexity of *O*(*r*) or *O*(*r*^rev^). In practice, however, the number of steps in the move structure is very small, as demonstrated by b-move’s results [22, 23], and the linear, cache-friendly memory access patterns make character extensions highly efficient. Consequently, these lossless run-length compressed move tables serve as a robust foundation for developing our new SMEM-finding and metagenomic classification implementation.

### 2.2 Finding SMEMs of length at least *L*

Before we can classify a read, we first find all of its *full* super-maximal exact matches (SMEMs) relative to the concatenation of all candidate sequences. Formally, a non-empty substring *P* [*i, j*) of a pattern *P* is an SMEM if it satisfies the following conditions: *P* [*i, j*) ∈ *T*, *P* [*i* − 1, *j*) ∉ *T* (or *i* = 0), and *P* [*i, j* + 1) ∉ *T* (or *j* = |*P*|). While some sources refer to these as MEMs [29], the term MEM is sometimes also used to describe a match that cannot be extended at a specific position in *T* but may still be extendable at other positions. For clarity, we continue using the term SMEMs.

#### Algorithm 1

Given a pattern *P*, a start (or end) position of interest, and the move table *M* ^rev^ (or *M*), this algorithm finds the corresponding SMEM’s end (or start) position using forward (or backward) pattern matching.

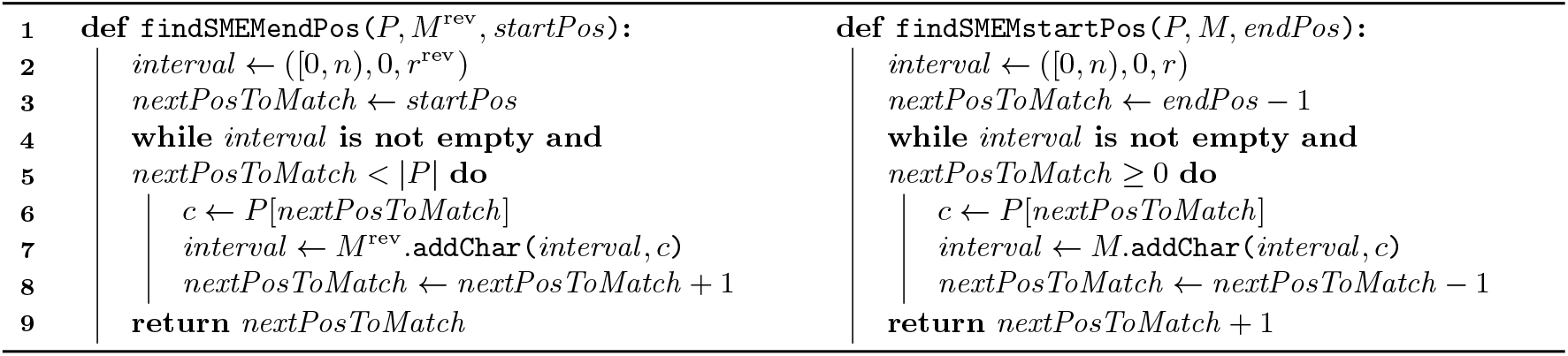

#### Algorithm 2

This algorithm finds all SMEMs of length at least *L* between pattern *P* and search text *T*, using its move tables *M* and *M* ^rev^.

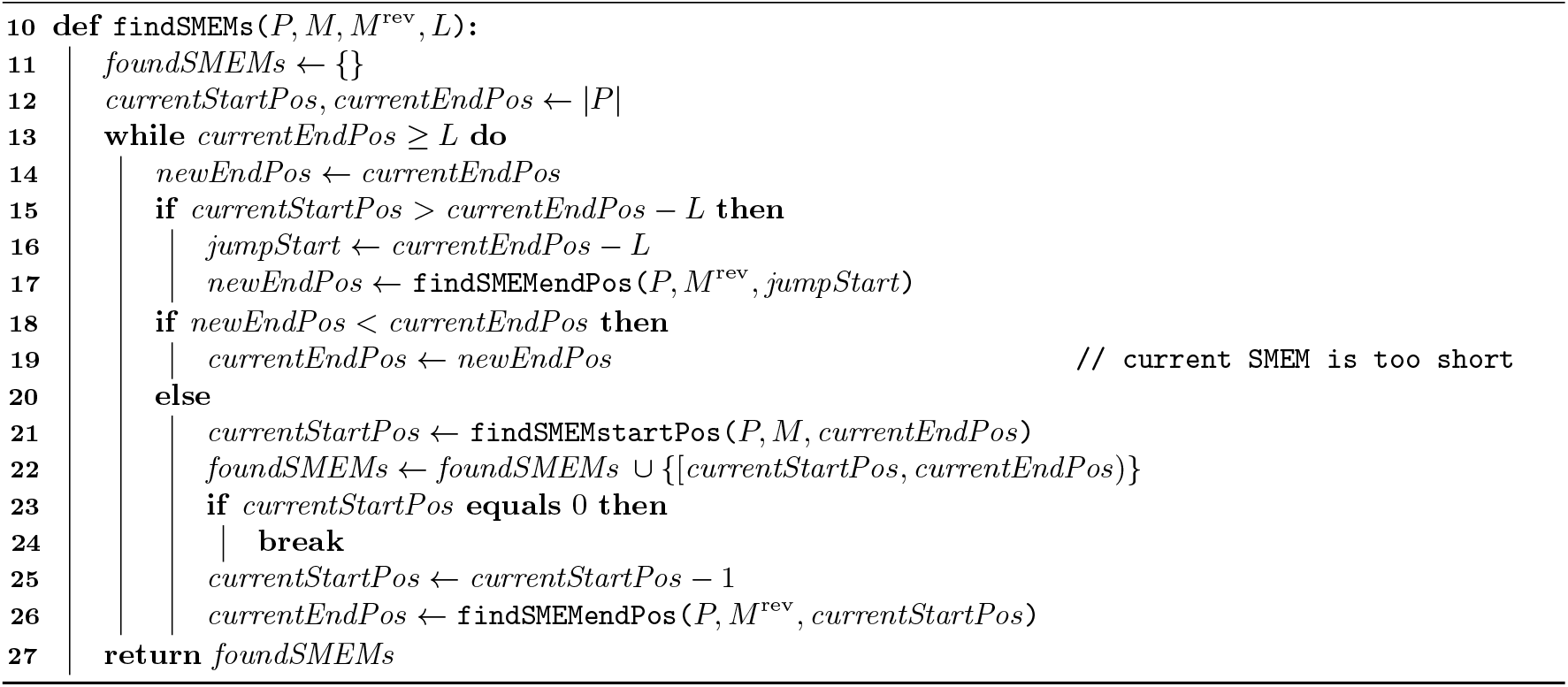

To identify all SMEMs between a pattern *P* and search text *T*, one can use the forward-backward search algorithm originally proposed by Li [28]. However, a substantial amount of computational time is spent identifying relatively short SMEMs between *P* and *T*. This issue is further exacerbated by the increasing adoption of large pan-genome references, which raise the likelihood that short substrings of *P* occur frequently in *T*. To address this, Gagie [29] proposed an updated forward-backward algorithm that enables skipping all SMEMs shorter than a customizable threshold *L*, which is also used in ropebwt3 [30]. This approach allows for a more efficient focus on SMEMs that are longer than expected by chance. In this section, we describe our implementation of a customized version of this parametrized forward-backward algorithm, leveraging solely the two move tables *M* and *M* ^rev^ introduced in Section 2.1.1.

Auxiliary Algorithm 1 and Algorithm 2 outline our approach to finding all SMEMs of length at least *L*. On line 12, we start at the end of pattern *P*. If the condition on line 15 fails (see further), the algorithm skips *L* characters ahead (in our case, backward) on line 16, which we refer to as a *jump start*. If the SMEM ending at the current *end* position is at least *L* characters long, it will bypass this jump start index. We can verify this by performing forward matching from the jump start toward the current end (line 17), using the auxiliary function in Algorithm 1 (which in turn calls the `addChar` functionality from Section 2.1.1). If we are unable to reach the current end position, the SMEM is shorter than *L*, and we skip it, starting with the new end position found during forward matching (line 19).

If, however, we do reach the correct end position, we know that an SMEM of length at least *L* ending at this position exists. We then compute the length of this SMEM by backward matching from this end position (line 21) and report the result on line 22. If the beginning of the pattern *P* has not yet been reached (line 23), we update the current end position by performing forward matching from the position immediately preceding the last found SMEM (line 26). The process then repeats. Note that if the part of the next SMEM that was found on line 26 is already of length at least *L*, the next jump start can be skipped (see line 15).

#### Example

We demonstrate the SMEM-finding algorithm on *P* = “CTATGTTGCTC”, a recombination of the example strains from Section 2.1.1 (“CTATGTC” and “ATATGTTGGTC”) with a “sequencing error” at the third-to-last position. For the sake of the example, we search for all SMEMs of length at least *L* = 3. Figure 1 shows the required character extensions, their order, and direction.

**Figure 1:**
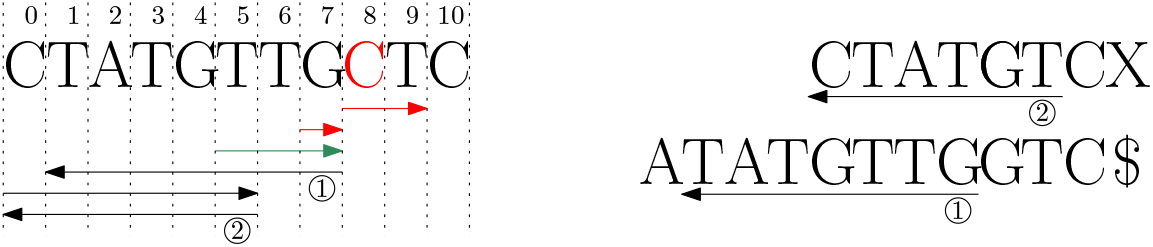
Left: visualization of the search steps in Algorithm 2 to execute `findSMEMs(`*P, M, M* ^rev^, 3), where *P* =“CTATGTTGCTC”, and *M* and *M* ^rev^ correspond to the example move tables in Table 1. *P* [8] = *C* represents a sequencing error. The red arrows (top two) represent failed jump start searches, the green arrow (third from the top) represents a successful jump start search. Black arrows (bottom three) represent searches that characterize valid SMEMs of length at least *L* = 3. Right: visualization of the detected SMEMs in the search text *T* .

The algorithm always starts with a jump start at the end of *P*. Due to the sequencing error at position 8, the first two jump starts (line 17) fail to match the 3-mers “CTC” and “GCT” in the search text. The third jump start succeeds, after which we locate the SMEM start position using a forward search (line 21). Subsequently, no further jump starts are needed, as the overlap with the next (and last) SMEM exceeds *L*− 1. This yields two SMEMs: *P* [1, 8) and *P* [0, 6), effectively skipping shorter SMEMs at the pattern’s end. Note that when a jump start succeeds, it performs some redundant work, as the subsequent call to `findSMEMstartPos` repeats the same character extensions. However, in practice, this redundancy is outweighed by the computational savings achieved through skipping short SMEMs with failed jump starts.

#### Lookup optimization

In Algorithms 1 and 2, a significant amount of time is spent matching the first characters of a partial match against the dense part of the search tree. To mitigate this, a lookup table can store forward and backward intervals for all possible *k*-mers, where *k* is set to 10 by default. Both intervals can be precomputed simultaneously using b-move’s bidirectional matching functionality [22, 23].

For the jump start on line 17, the match length is required to be *L*, so the lookup table can be applied assuming that *L*≥ *k*. If the lookup fails, we lose precision over the new *currentEndPos* value, but the performance gains outweigh this limitation in practice. Similarly, on line 21, we know the SMEM length is at least *L*≥ *k*, allowing use of the lookup table. Only the call to `findSMEMendPos` on line 26 lacks guarantees on match length, so the lookup table cannot be applied there.

Including the lookup table increases the index’s theoretical space complexity from *O*(*r* + *r*^rev^) to *O*(*r* + *r*^rev^ + 4^*k*^), assuming *L* ≥ *k*. However, in practical use cases, the lookup table’s space usage remains relatively small compared to the other index components.

### 2.3 Assigning a class tag to each SMEM

#### 2.3.1 Storing a sampled tag array

To perform metagenomic read classification, we assign a metagenomic class identifier to each SMEM. To do this, we store a sampled tag array alongside the two move tables. A tag array “tags” each position in the BWT with metadata [31], such as positions in a pan-genome graph [32] or, in our case, metagenomic class identifiers. In our tag array, each position in the BWT is tagged with its corresponding metagenomic class identifier. Realistic datasets typically contain multiple species within each class, so sequences from the same class tend to cluster together in the BWT. While this contextual locality could be used to directly run-length compress the tag array [31], we opt to sample the tag array at the BWT run end positions. Sampling at the BWT run ends retains *O*(*r* + *r*^rev^) memory complexity and proves more memory-efficient for datasets with many metagenomic classes, without negatively impacting classification performance.

##### Algorithm 3

Update the tag toehold for a partial match *P* with interval ([*s, e*), *R*_*s*_, *R*_*e*_), using move table *M* and sampled tag array *t*. The match *P*, corresponding to *previousTagToehold*, will be extended by prepending character *c*.

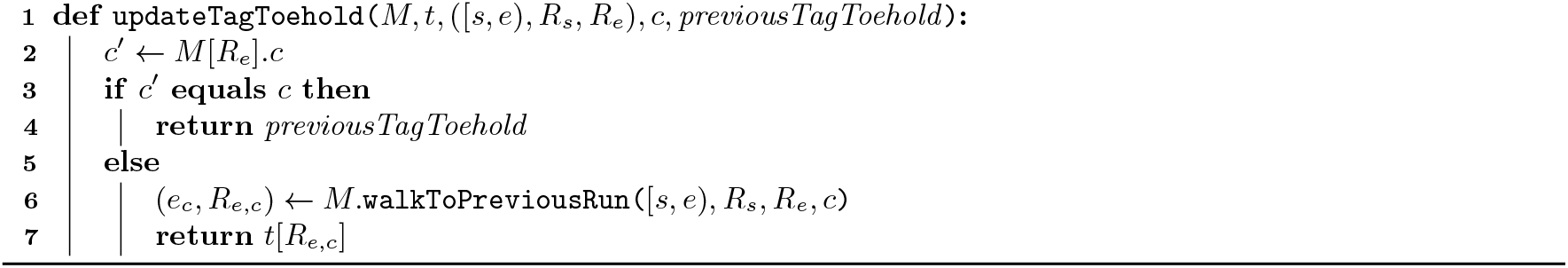

##### Example

In our example, let the first strain (“CTATGTC”) correspond to metagenomic class 0 and the second strain (“ATATGTTGGTC”) to class 1. Table 2 shows the corresponding sampled tag array *t*. Dashes indicate positions that can contain any value, as they are never queried (they correspond to prefixes starting with “$” or “X”). For completeness, the unsampled tag array is provided in Supplementary Table S1. Note that this example is too small to demonstrate contextual locality, as it contains only one sequence per class.

**Table 2:** The sampled tag array *t*, containing *r* metagenomic class identifiers, each corresponding to a run in the move table *M* (see Table 1). The full tag array is provided in Supplementary Table S1.

| $j$ | 0 | 1 | 2 | 3 | 4 | 5 | 6 | 7 | 8 | 9 | 10 | 11 | 12 | 13 |
| --- | --- | --- | --- | --- | --- | --- | --- | --- | --- | --- | --- | --- | --- | --- |
| $t[j]$ | - | 1 | 1 | 0 | 1 | 1 | 1 | 0 | 1 | 0 | 1 | 1 | 1 | - |

#### 2.3.2 Updating the tag toehold

To assign a class tag to each SMEM, we track *one* tag value while matching the SMEM to the index. We call this value the “tag toehold”, analogous to the suffix array toehold kept in the r-index [15, 16]. This tag toehold always represents the class tag of the *last* index within the BWT interval of the current partial match. When the full SMEM is identified, it is assigned that tag of its last occurrence in the BWT, even if it appears in other metagenomic classes. Since we search for SMEMs of length at least *L* (i.e., with high specificity) and expect contextual locality in the tag array, this single tag is assumed to be representative in most cases. This approach avoids the computationally expensive process of querying the class tags for *all* occurrences of an SMEM, which can be numerous in pan-genomic contexts.

To maintain the tag toehold while backward matching an SMEM (line 21, Alg. 2), it must be updated each time *before* a successful `addChar` operation is performed in `findSMEMstartPos` (Alg. 1). Algorithm 3 outlines this process. The update starts with the BWT interval [*s, e*) of the current partial match *P*, the run indices *R*_*s*_ and *R*_*e*_ corresponding to BWT indices *s* and *e*− 1, the character *c* about to be prepended to *P*, and the tag toehold from the previous iteration. For the first iteration, the tag toehold is initialized to *t*[*r*− 1]. The algorithm checks if the character *c*^*′*^ of BWT run *R*_*e*_ matches *c* on line 3. If so, the last occurrence of *P* ’s BWT interval also corresponds to the last occurrence of *cP* ’s BWT interval, and the previous tag toehold remains valid (line 4). Otherwise, the largest BWT index (*e*_*c*_ − 1) ∈ [*s, e*) with BWT[*e*_*c*_ − 1] = *c* and its run index *R*_*e,c*_ are located using the `walkToPreviousRun` function from Section 2.1.1 (line 6). The updated tag toehold is then set to the last tag of run *R*_*e,c*_, which is sampled at *t*[*R*_*e,c*_]. If no such *e*_*c*_ index exists, the end of the SMEM has been found and the toehold must no longer be updated.

##### Example

We illustrate the toehold updating process for SMEM *P* [0, 6) = “CTATGT”, found for pattern *P* = “CTATGTTGCTC” in Section 2.2. All operations can be verified using Tables 1 and S1. Starting with a lookup table where *k* = 3, the interval for *P* [3, 6) = “TGT” is ([16, 18), 11, 11), with tag toehold 1. The run character *M* [11].*c* = “A” matches the next character of *P*, so the toehold remains 1. After extending with “A”, the interval for *P* [2, 6) = “ATGT” becomes ([2, 4), 2, 2). Again, *M* [2].*c* = “T” matches the next character in *P*, so the toehold remains 1. Next, the interval for *P* [1, 6) = “TATGT” is ([11, 13), 7, 8). Here, *M* [8].*c* = “A” does not match *P* [0] = “C”, so we compute *M*.`walkToPreviousRun(`[11, 13), 7, 8, “C”) = (12, 7), where 12 is the exclusive upper bound, yielding final toehold *t*[7] = 0.

### 2.4 Consensus classification

After processing a pattern and identifying its SMEMs with tag toeholds, we assign a consensus metagenomic class to the read. This is done by tracking the accumulated SMEM length for each class as evidence. To avoid bias, overlapping SMEM regions from the same class are counted only once. This ensures that multiple shorter, overlapping SMEMs do not outweigh a single long, high-evidence SMEM. The class with the highest evidence from non-overlapping SMEMs is selected as the consensus tag. Ties are broken randomly.

For read data, the process is repeated for the reverse complement. The final consensus tag is based on the orientation (forward or reverse complement) with the most evidence across all classes.

## 3 Results

### 3.1 Setup and hardware

We compare the accuracy and performance of our metagenomic read classification implementation with SPUMONI 2 [24] and Cliffy [26], the only run-length compressed tools, to our knowledge, supporting multi-class metagenomic read classification without relying on taxonomy. SPUMONI 2 can build and query indexes with and without minimizer digestion^1^. Minimizer digestion reduces runtime and memory usage, but introduces a slight accuracy penalty due to the use of a lossy index. Both configurations are evaluated. SPUMONI 2 produces pseudomatching lengths, each with *one* corresponding class identifier, for each position in the read *P*. From these |*P*| identifiers, we determine the consensus class using a majority vote, as described in the SPUMONI 2 paper [24]. Post-processing majority vote runtimes are excluded from benchmarks.

Cliffy, though mainly intended for taxonomic classification, also supports multi-class metagenomic classification^2^. Cliffy offers configurations with or without minimizer digestion, with similar trade-offs to SPUMONI 2. Orthogonally, Cliffy supports uncompressed and cliff compressed document array profiles. Uncompressed profiles enable *exact* document listing but are practical only for datasets with few metagenomic classes. To support many metagenomic classes, cliff compression is required to maintain a manageable index size, resulting in *approximate* document listing. Depending on the dataset, we will use the Cliffy configurations that result in a reasonably-sized index. For each read, Cliffy produces variable-length exact matches, each with a set of candidate classes. We assign a single metagenomic class to each read using a consensus algorithm that weighs candidate classes by match length and the number of classes per match, following the method by Ahmed et al. [26]. This consensus algorithm is excluded from runtime benchmarks.

All benchmarks used a single core of two 18-core Intel® Xeon® Gold 6140 CPUs (2.30 GHz) and 177 GiB of RAM. Runtimes represent the median of 20 runs and exclude index loading times.

### 3.2 Case study with 8 165 bacterial genomes

We performed a case study similar to the SPUMONI 2 multi-class classification experiment [24]. A pangenome was created by combining all RefSeq strains from eight bacterial species: *Escherichia coli, Salmonella enterica, Listeria monocytogenes, Pseudomonas aeruginosa, Bacillus subtilis, Limosilactobacillus fermentum, Enterococcus faecalis*, and *Staphylococcus aureus*. This pan-genome contains 8 165 sequences^3^ across 8 metagenomic classes, totaling 37.4 Gbp. To assess read classification accuracy, we simulated 50 000 long reads^4^ [33] (average length 5 236 bp) and 500 000 short reads^5^ [34] (length 250 bp) per species.

Figure 2 compares the accuracy and runtime performance of our approach against SPUMONI 2 and Cliffy for this dataset (an extended version of the bottom left plot is shown in Supplementary Figure S1). For Cliffy, only the index with minimizer digests is included, as its counterpart without minimizers exceeds our workstation’s RAM capacity. Both uncompressed and cliff compressed document array profiles can be analyzed due to the small number of metagenomic classes. Our approach (for optimal *L*) outperforms SPUMONI 2 and Cliffy in accuracy across all configurations, as shown in the left panels. While the improvement over SPUMONI 2 is modest, especially for short reads, Cliffy demonstrates noticeably lower accuracy overall. Specifically, cliff compressed document array profiles show a significant reduction in accuracy for this dataset, particularly for long reads. This indicates that cliff compression is not suited for datasets with few metagenomic classes.

**Figure 2:**
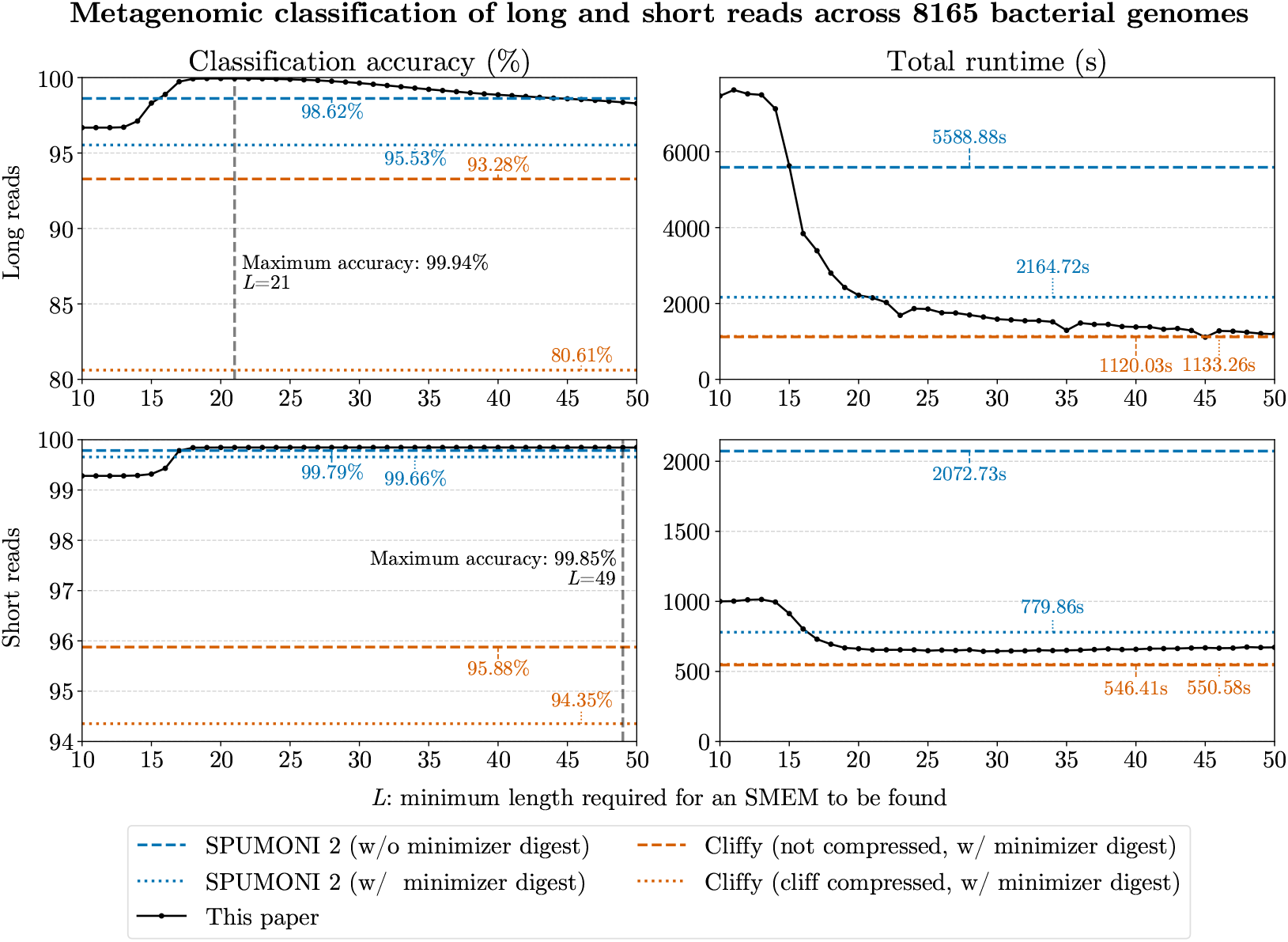
Analysis of metagenomic classification accuracy (left) and runtime performance (right) for matching 400 000 long reads (top) and 4 000 000 short reads (bottom) to 8 165 bacterial genomes across 8 metagenomic classes. We compare our approach for varying values of *L*, to SPUMONI 2 (without/with minimizer digests) and Cliffy with minimizer digests (with uncompressed/cliff compressed document array profiles).

In terms of runtime (right panels), our approach outperforms SPUMONI 2 when *L* is sufficiently large (*L* ≥ 21 for long reads and SPUMONI 2 using minimizer digests, or even smaller for other cases). Conveniently, this value of *L* aligns with our best classification accuracy (also for short reads, accuracy is 99.85% at *L* = 21). We are unable to beat Cliffy’s runtime; however, given Cliffy’s significantly lower accuracy, runtime comparisons are less meaningful.

#### Per-class analysis

To assess potential bias, such as towards more abundant classes, we analyze the accuracy, sensitivity, specificity, and precision per class for the long read experiment with optimal parameter *L* = 21 (see Supplementary Figure S2). Our results show that the approach performs consistently across all 8 species in the classification task, regardless of abundance. The most noticeable deviations are lower precision for *E. coli* and lower sensitivity for *S. enterica*, which arise from 161 *S. enterica* reads being misclassified as *E. coli*. Since *S. enterica* is closely related to *E. coli* but possesses a significant number of additional virulence genes [35], these misclassifications are not unexpected.

#### SMEM length distribution

To demonstrate the benefit of restricting the SMEM search to a minimum length of *L*, we analyze the distribution of the lengths of all SMEMs found during the long read experiment for *L* = 1 (see Supplementary Figure S3). We also visualize the fraction of these SMEMs that have a correct tag toehold with respect to the read they originate from. We observe a significant peak in SMEMs shorter than 20 bases, with the highest frequency at length 14. Moreover, most of these short SMEMs correspond to incorrect tag toeholds. This confirms that, beyond reducing runtime, setting a minimum SMEM length of *L* ≥ 20 improves SMEM precision.

### 3.3 Case study with 402 378 16S rRNA genes

To demonstrate the versatility of our approach, we also evaluated its performance on the SILVA SSU NR99 (version 138.1) database, which contains 510 508 16S rRNA sequences, similar to Cliffy’s analysis [26]. From this collection, we selected the 402 378 16S rRNA gene sequences^6^ annotated to the genus level, totaling to 586 Mbp of data spanning 9 118 genera, which serve as the metagenomic classes. Compared to the dataset in Section 3.2, this dataset is smaller in size but contains significantly more metagenomic classes. We simulated 5 726 966 long reads (average length 1 415 bp) and 9 029 669 short reads (length 250 bp), aiming for equal coverage across genera.

Figure 3 compares our approach to SPUMONI 2 and Cliffy (an extended version of the bottom left plot is provided in Supplementary Figure S4). For the SILVA dataset, only Cliffy indexes with cliff compressed document array profiles are considered, as their uncompressed counterparts exceed 1 TiB of memory [26]. For classification accuracy, our approach (for optimal *L*) consistently outperforms SPUMONI 2 for both read types. However, Cliffy without minimizers achieves the highest accuracy across all tools. For long reads, Cliffy with minimizers also outperforms our method. This suggests that for datasets with many similar metagenomic classes, Cliffy’s ability to report multiple classes per match provides a significant advantage.

**Figure 3:**
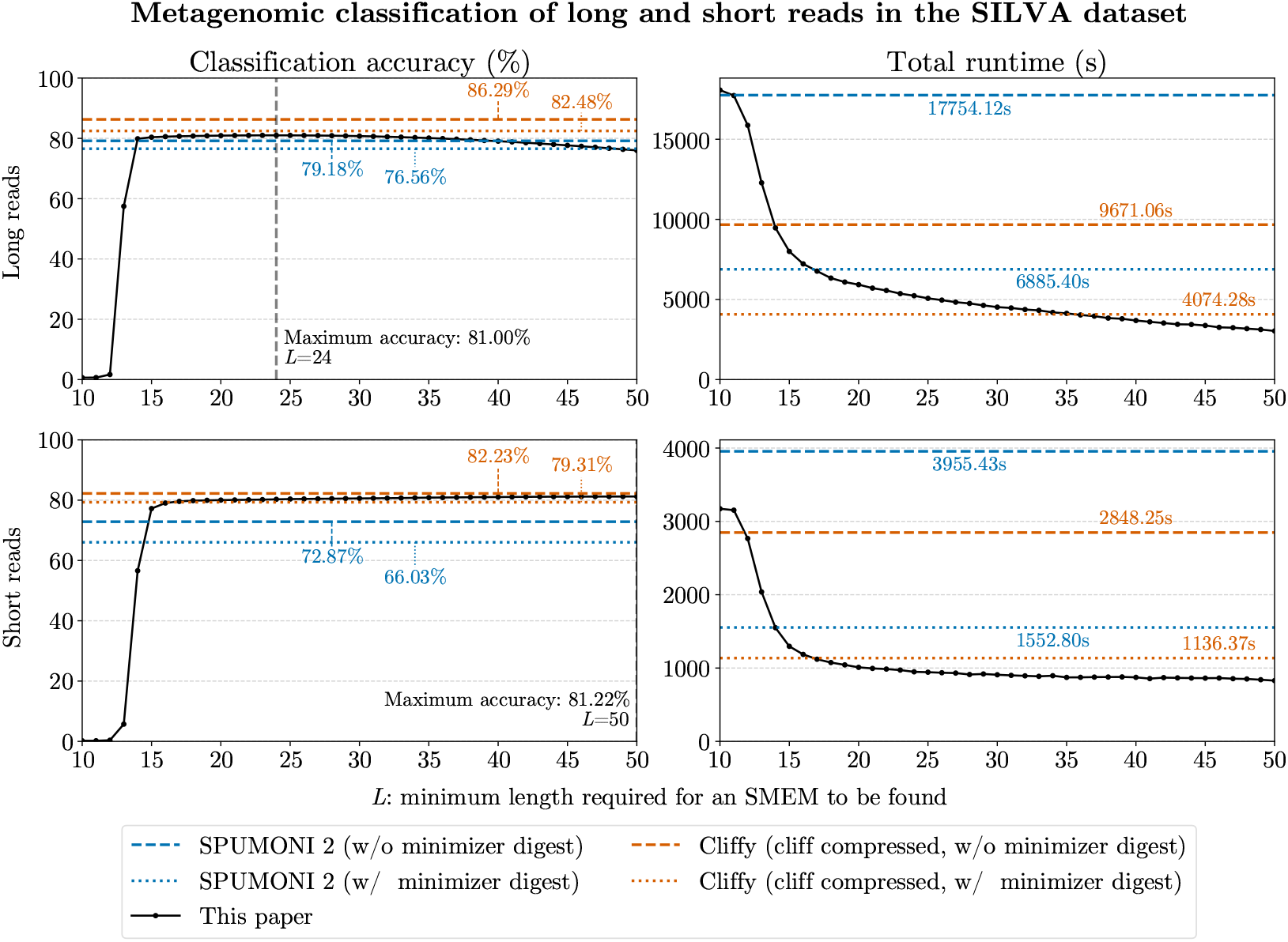
Analysis of metagenomic classification accuracy (left) and runtime performance (right) for matching 5 726 966 long reads (top) and 9 029 669 short reads (bottom) to 402 378 16S rRNA gene sequences spanning 9 118 metagenomic classes. We compare our approach for varying values of *L*, to SPUMONI 2 (without/with minimizer digests) and Cliffy with cliff compressed document array profiles (without/with minimizer digests).

In terms of runtime, our method again significantly outperforms SPUMONI 2 across all configurations. For this dataset, we also surpass Cliffy in runtime, except when matching long reads with *L* ≤35, where Cliffy with minimizer digests achieves both higher accuracy and faster performance.

### 3.4 Memory versus runtime analysis

Finally, we compare the tools in terms of peak memory usage and runtime performance. Figure 4 shows memory versus runtime for all long read experiments (Sections 3.2 and 3.3), and Supplementary Figure S5 covers short reads. For our approach, we fix *L* at 25, balancing accuracy and runtime as demonstrated above. For the bacterial genomes dataset, where Cliffy shows significantly lower accuracy (Section 3.2), Cliffy (uncompressed and cliff compressed) uses 2.4× and 11.5× more memory than our method, respectively. In combination with Cliffy’s poor accuracy on this dataset, particularly with cliff compression, these results confirm that Cliffy is less suitable for larger datasets with few metagenomic classes, especially when cliff compression is involved. Compared to SPUMONI 2 (without and with minimizer digests), our method uses 4.1× and 8.7 ×more memory, respectively, but is faster and more accurate. Our index, under 8 GiB, still fits within standard laptop RAM.

**Figure 4:**
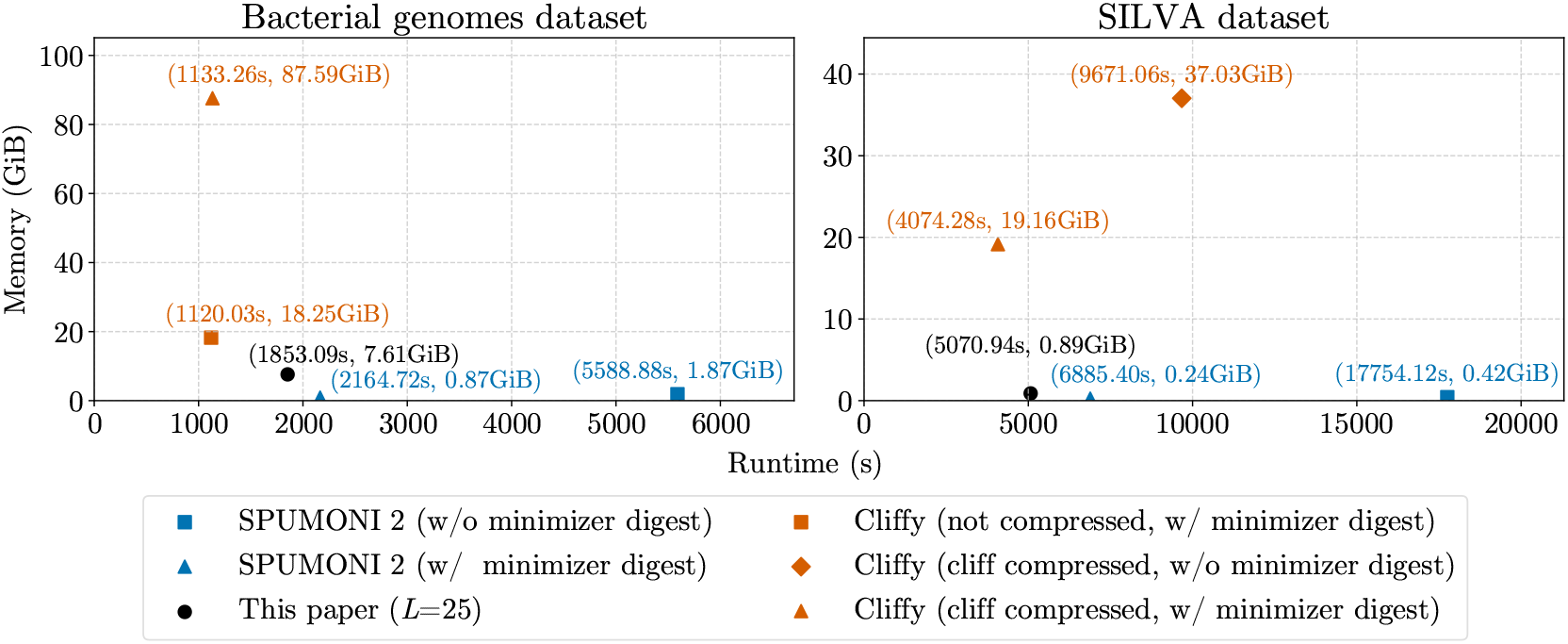
Memory versus runtime comparison of our approach (*L* = 25) with SPUMONI 2 and Cliffy, using all configurations the dataset allows. Shown are results for matching 400 000 long reads to an index of 8 165 bacterial genomes across 8 metagenomic classes (left); and for matching 5 726 966 long reads to an index of 402 378 16S rRNA gene sequences spanning 9 118 metagenomic classes (right).

For the SILVA rRNA dataset, the memory difference between Cliffy and the other tools is even more pronounced. Cliffy requires 41.8× and 21.6× more memory than our method (without and with minimizer digests, respectively). Cliffy’s higher accuracy (Section 3.3) thus comes at the cost of large document array profiles. For instance, Cliffy (with minimizer digests) is more accurate and faster in one case (Figure 3, top right) but requires nearly 20 GiB of RAM, while our approach uses less than 1 GiB. This could affect the feasibility of running Cliffy on a portable laptop. Compared to SPUMONI 2 (without and with minimizer digests), our method requires 2.2 ×and 3.7× more memory, respectively, but is again faster and more accurate. Both our index and SPUMONI 2’s indexes fit within 1 GiB, however.

## 4 Discussion

### Index construction

Since the index is constructed only once, less time was spent optimizing this part of the implementation. We support two types of index construction: one based on the full suffix array and another using prefix-free parsing [36]. The suffix array approach is faster but memory-intensive, suitable for smaller datasets like the SILVA dataset (under 1 hour runtime, 6.2 GiB peak RAM). Prefix-free parsing is slower but uses less memory, making it ideal for larger pan-genomes like the bacterial genomes dataset (approximately 13 hours runtime, 106 GiB peak RAM, feasible on a regular workstation). We aim to improve our construction implementation based on recent advancements such as grlBWT [37] and ropebwt3 [30].

### Alternate or additional implementations

In addition to the algorithms presented above, we explored several alternate implementations, details of which are beyond the scope of this paper, that were outperformed by our current approach. One alternative sampled the tag array at the ends of tag runs instead of BWT runs, using a sparse bit vector to mark these sampled positions. This bit vector could identify SMEMs where all occurrences correspond to the same metagenomic class. Another modification combined this bit vector with the tag array sampled at BWT run ends to retain information on SMEMs with a unique class identifier. However, including information on unique SMEMs did not improve accuracy and was thus omitted. We also considered full read alignment for short reads [38] instead of finding SMEMs, but this did not yield better performance or accuracy.

Additionally, we support subsampling the tag array to reduce its memory usage, at the cost of a slight runtime performance penalty, similar to subsampling the suffix array samples in the sr-index [39, 40]. However, the relative memory reduction is modest, as the majority of the index consists of the two move tables. Finally, we also support parallelization.

### Future work

We plan to optimize and build our index for large, taxonomically comprehensive datasets such as The Genome Taxonomy Database (GTDB) [41, 42] and AllTheBacteria [43], making prebuilt versions publicly available. This would allow researchers to classify metagenomic reads efficiently without requiring expensive index construction. Additionally, we aim to integrate phylogenetic tree information within the index to enable more advanced taxonomic classification, further enhancing our index’s practical utility in metagenomic workflows.

## 5 Conclusion

We introduced a novel run-length compressed approach to metagenomic read classification based on the move structure, SMEM-finding, and sampled tag arrays. Our method demonstrates versatility, achieving high accuracy across various read types (long and short) and datasets, including a large pan-genome with few metagenomic classes and a smaller rRNA database with thousands of classes. We consistently outperform SPUMONI 2, being both faster and more accurate, while maintaining the same memory complexity of *O*(*r*). While Cliffy achieves slightly higher accuracy for the complex dataset with thousands of metagenomic classes, this comes at the cost of a significant memory increase (20 to 40 times more). Thus, our approach offers a balanced solution, demonstrating strong performance in accuracy, runtime performance, and memory efficiency, making it suitable for a wide range of metagenomic classification tasks. Our open-source C++11 implementation is available at https://github.com/biointec/tagger under the AGPL-3.0 license.

## Acknowledgements

Lore Depuydt was funded by a PhD Fellowship FR (1117322N), Research Foundation – Flanders (FWO). Travis Gagie was funded by NSERC grant 07185-2020. Ben Langmead and Omar Ahmed were funded by NSF grant DBI-2029552 and NIH grant R01HG011392.

---

**Table S1:**
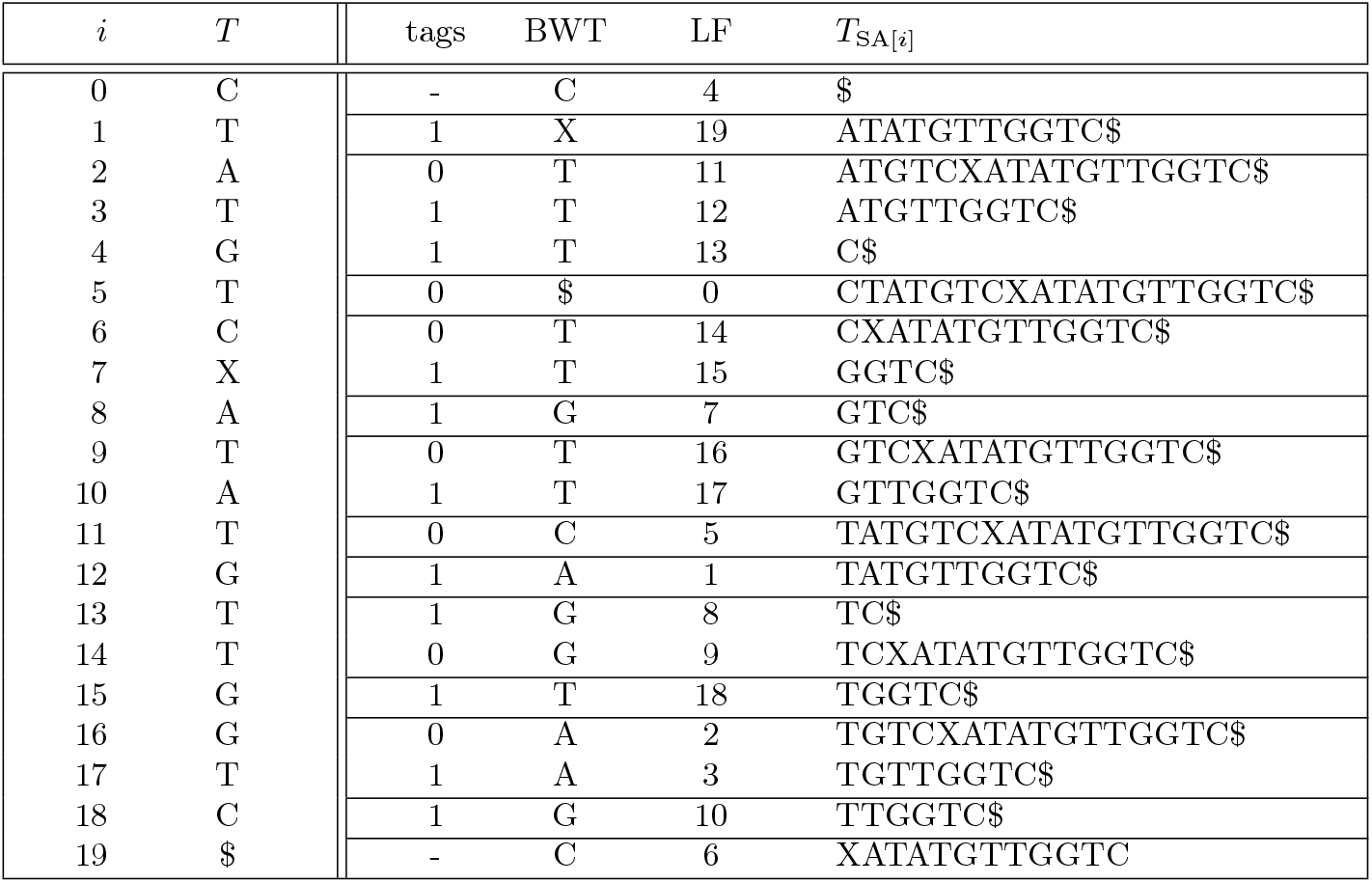
FM-index for the search text *T* = “CTATGTCXATATGTTGGTC$” showing its tag array, Burrows-Wheeler transform BWT (Last column), LF mapping, and suffixes (with the First column represented by the suffixes’ first characters). BWT runs are separated by horizontal lines.

**Table S2:** FM-index for the reverse search text *T* ^rev^ = “$CTGGTTGTATAXCTGTATC” showing its Burrows-Wheeler transform BWT^rev^, LF mapping LF^rev^, and suffixes. BWT runs are separated by horizontal lines.

| $i$ | $T^{\text{rev}}$ | $\text{BWT}^{\text{rev}}$ | $\text{LF}^{\text{rev}}$ | $T_{\text{SA}^{\text{rev}}[i]}^{\text{rev}}$ |
| --- | --- | --- | --- | --- |
| 0 | \$ | C | 4 | \$CTGGTTGTATAXCTGTATC |
| 1 | C | T | 11 | ATAXCTGTATC |
| 2 | T | T | 12 | ATC |
| 3 | G | T | 13 | AXCTGTATC |
| 4 | G | T | 14 | C |
| 5 | T | \$ | 0 | CTGGTTGTATAXCTGTATC |
| 6 | T | X | 19 | CTGTATC |
| 7 | G | T | 15 | GGTTGTATAXCTGTATC |
| 8 | T | T | 16 | GTATAXCTGTATC |
| 9 | A | T | 17 | GTATC |
| 10 | T | G | 7 | GTTGTATAXCTGTATC |
| 11 | A | G | 8 | TATAXCTGTATC |
| 12 | X | G | 9 | TATC |
| 13 | C | A | 1 | TAXCTGTATC |
| 14 | T | A | 2 | TC |
| 15 | G | C | 5 | TGGTTGTATAXCTGTATC |
| 16 | T | T | 18 | TGTATAXCTGTATC |
| 17 | A | C | 6 | TGTATC |
| 18 | T | G | 10 | TTGTATAXCTGTATC |
| 19 | C | A | 3 | XCTGTATC |

**Figure S1:**
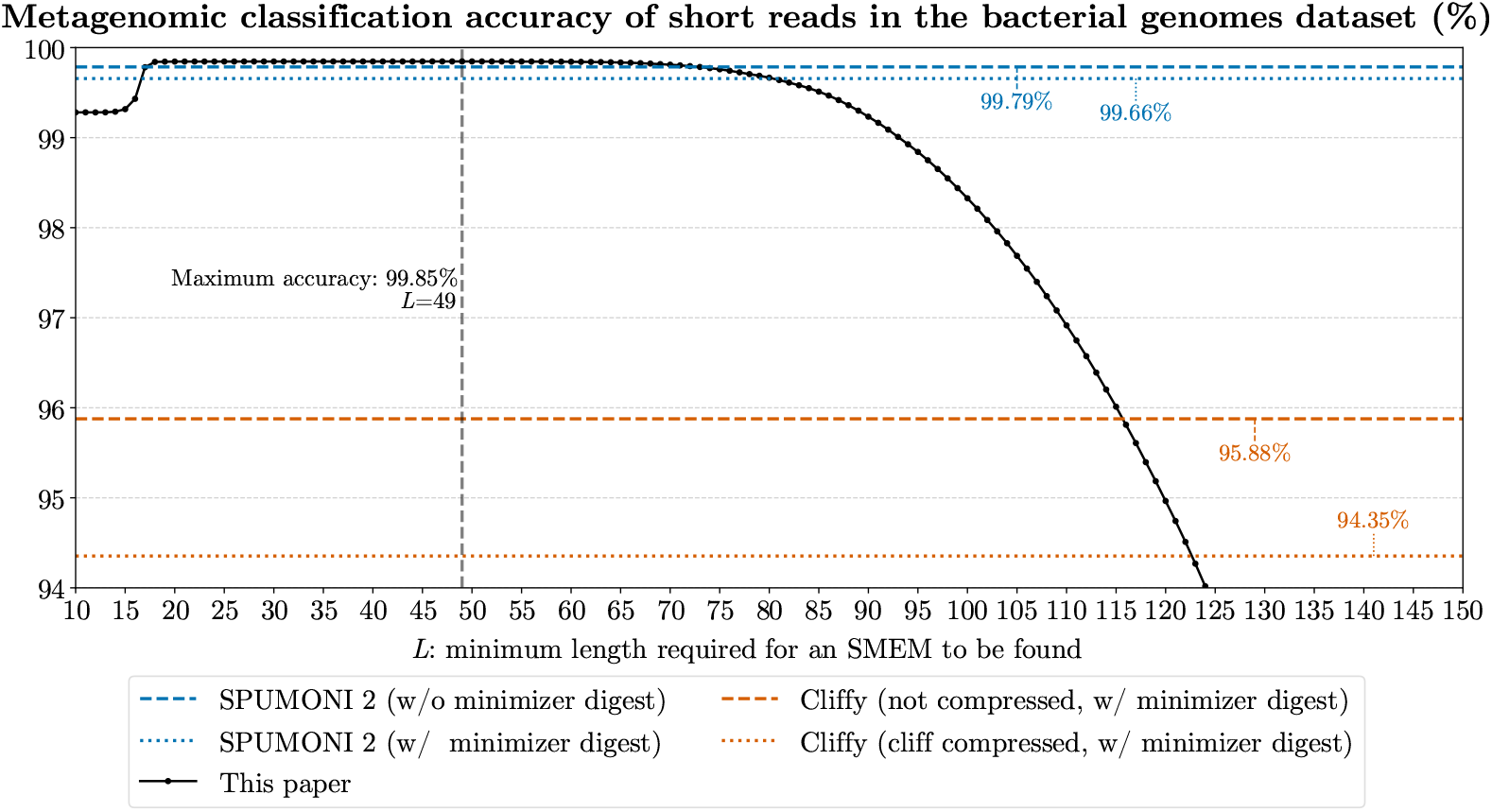
Analysis of metagenomic classification accuracy for matching 4 000 000 short reads to an index of 8 165 bacterial genomes across 8 metagenomic classes. We compare our approach for varying values of *L*, to SPUMONI 2 (without and with minimizer digests) and Cliffy with minimizer digests (with uncompressed and cliff compressed document array profiles). This is an extended version of the bottom left panel of Figure 2.

**Figure S2:**
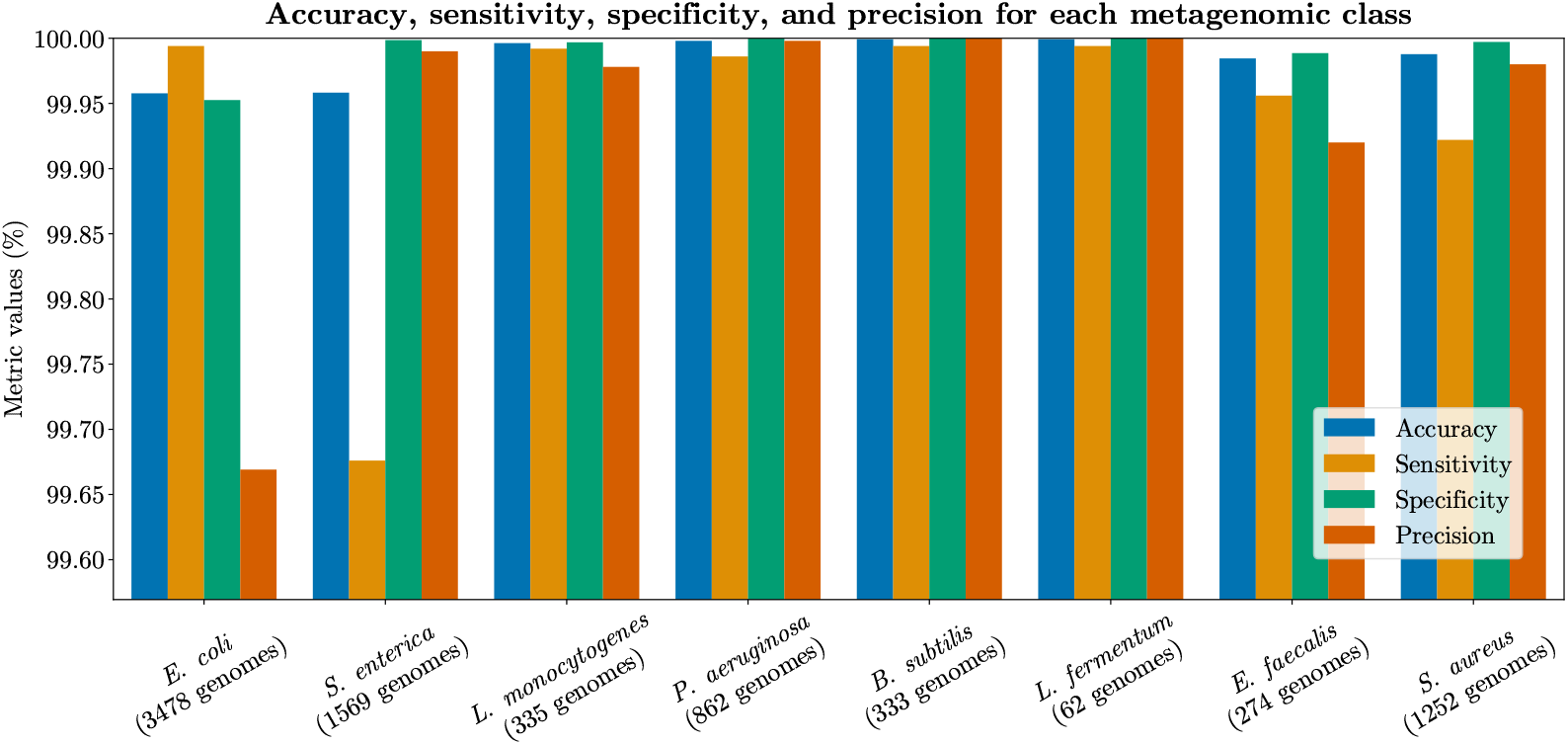
Per-class analysis of classification accuracy, sensitivity, specificity, and precision for matching 400 000 long reads to 8 165 bacterial genomes across 8 metagenomic classes (*L* = 21).

**Figure S3:**
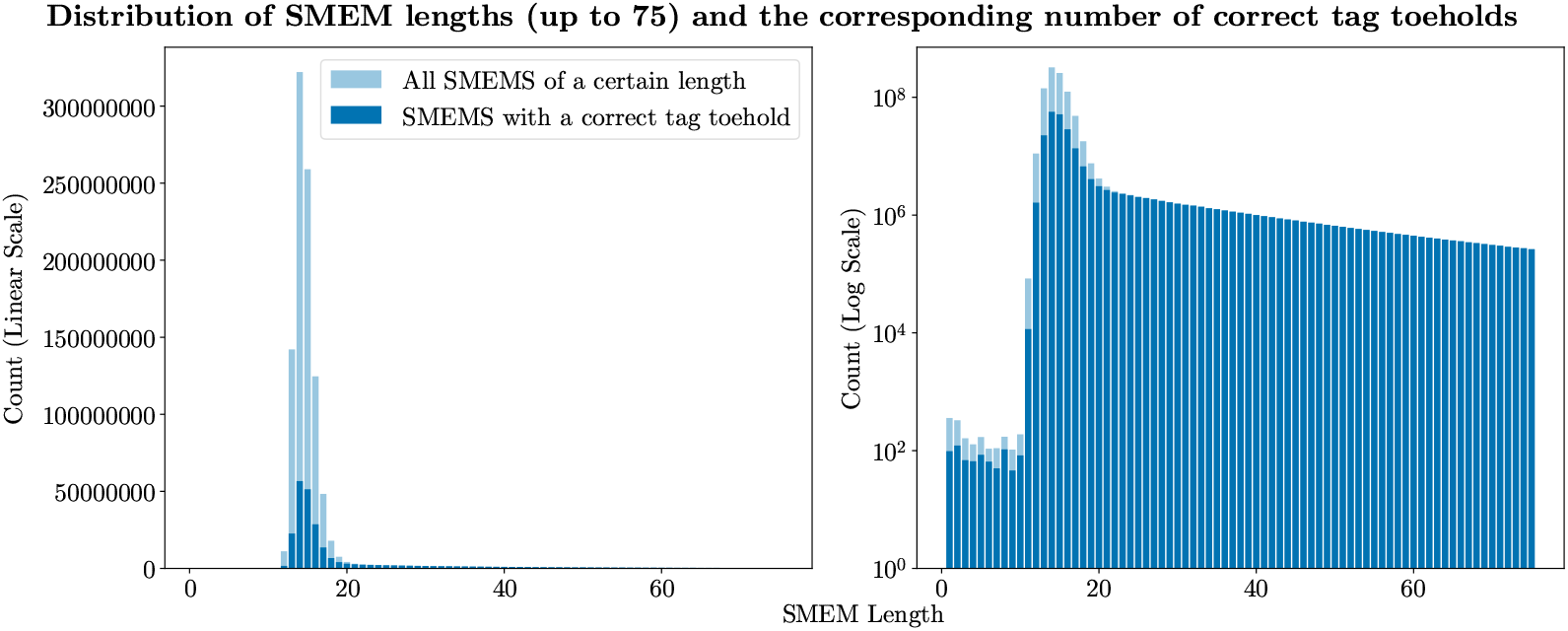
Distribution of SMEM lengths and the corresponding counts of correct tag toeholds for matching 400 000 long reads to 8 165 bacterial genomes across 8 metagenomic classes (with *L* = 1). The distribution is shown in both linear (left) and log (right) scales.

**Figure S4:**
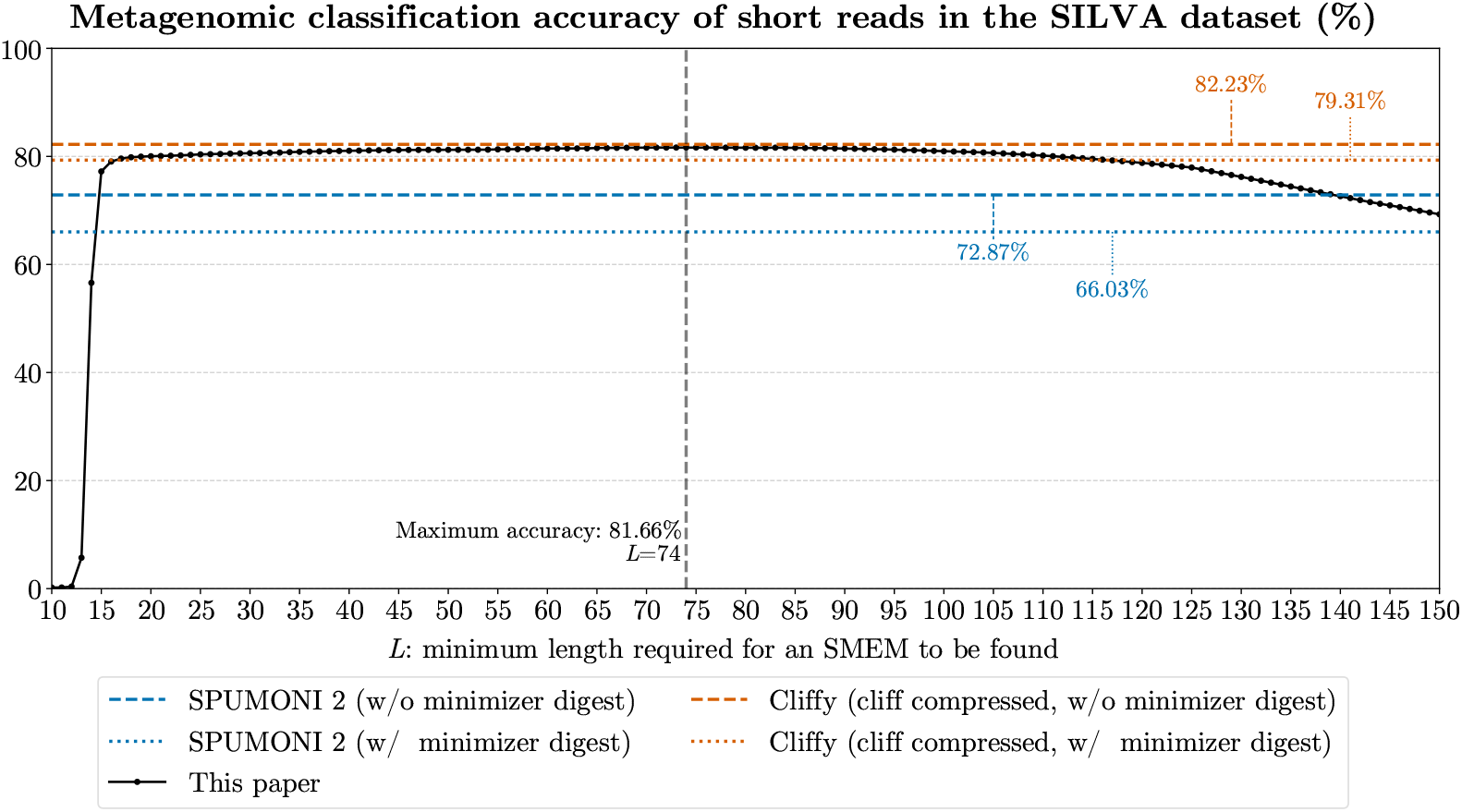
Analysis of metagenomic classification accuracy for matching 9 029 669 short reads to an index of 402 378 16S rRNA gene sequences spanning 9 118 metagenomic classes. We compare our approach for varying values of *L*, to SPUMONI 2 (without and with minimizer digests) and Cliffy with cliff compressed document array profiles (without and with minimizer digests). This is an extended version of the bottom left panel of Figure 3.

**Figure S5:**
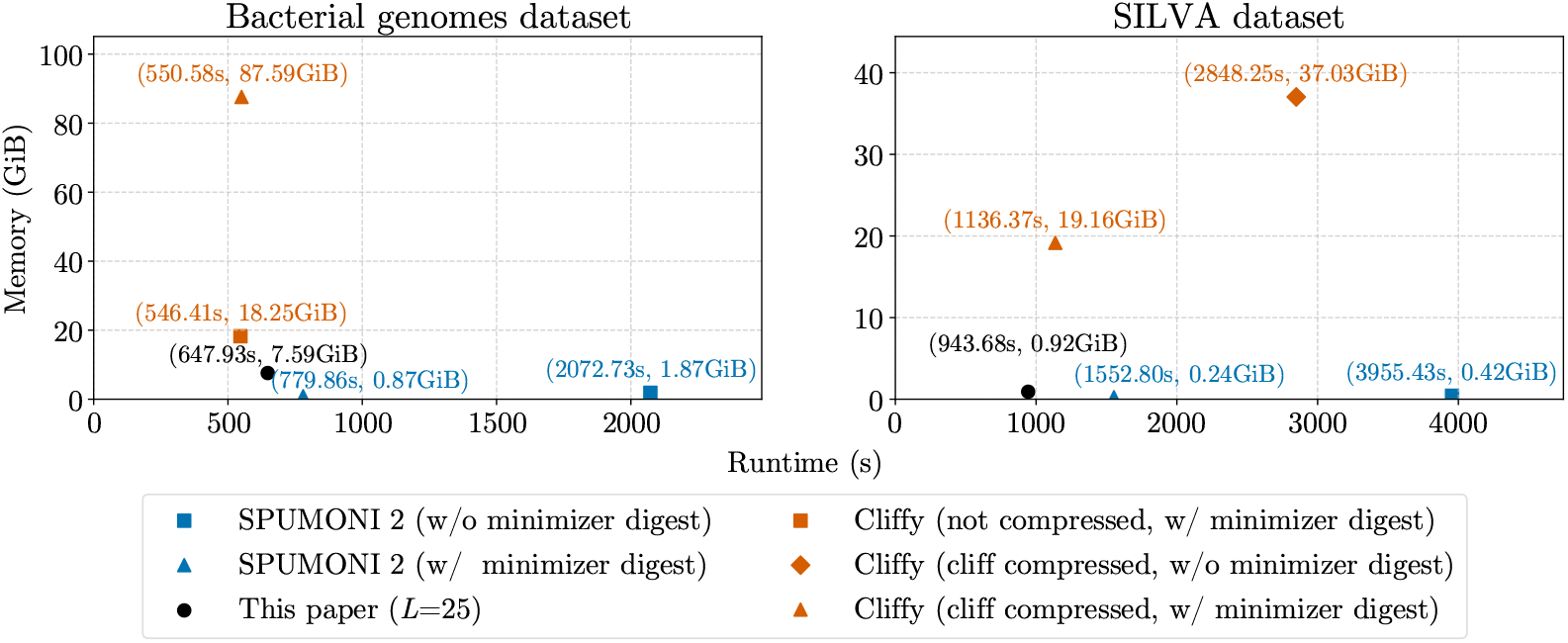
Memory versus runtime comparison of our approach (*L* = 25) with SPUMONI 2 and Cliffy, using all configurations the dataset allows. Shown are results for matching 4 000 000 short reads to an index of 8 165 bacterial genomes across 8 metagenomic classes (left); and for matching 9 029 669 short reads to an index of 402 378 16S rRNA gene sequences spanning 9 118 metagenomic classes (right).

## Footnotes

1 `SPUMONI v2.0.9: spumoni build/run --PML --doc-array --no-digest/--minimizer-alphabet`

2 `cliffy v2.0.0: cliffy build --revcomp --two-pass [--minimizers] [--taxcomp --num-col 7]; cliffy run --ftab [--minimizers] [--taxcomp --num-col 7]`

3 Details available at https://github.com/biointec/tagger/tree/data/BacterialGenomes

4 `PBSIM v2.0.1: pbsim --hmm model R94.model --accuracy-mean 0.95`

5 ART Illumina v2.3.7: art illumina --seqSys MS --len 250

6 Details available at https://github.com/biointec/tagger/tree/data/SILVA

